# Stable networks of water-mediated interactions are conserved in activation of diverse GPCRs

**DOI:** 10.1101/351502

**Authors:** A. J. Venkatakrishnan, Anthony K. Ma, Rasmus Fonseca, Naomi R. Latorraca, Brendan Kelly, Robin M. Betz, Chaitanya Asawa, Brian K. Kobilka, Dror Ron O.

**Affiliations:** Department of Computer Science, Stanford University, Stanford, California 94305, USA.; Institute for Computational and Mathematical Engineering, Stanford University, Stanford, California 94305, USA.; Department of Molecular and Cellular Physiology, Stanford University School of Medicine, Stanford, California 94305, USA.; Biophysics Program, Stanford University, Stanford, California 94305, USA.

## Abstract

G protein-coupled receptors (GPCRs) have evolved to recognize incredibly diverse extracellular ligands while sharing a common architecture and structurally conserved intracellular signaling partners. It remains unclear how binding of diverse ligands brings about GPCR activation, the common structural change that enables intracellular signaling. Here, we identify highly conserved networks of water-mediated interactions that play a central role in activation. Using atomic-level simulations of diverse GPCRs, we show that most of the water molecules in GPCR crystal structures are highly mobile. Several water molecules near the G protein-coupling interface, however, are stable. These water molecules form two kinds of polar networks that are conserved across diverse GPCRs: (i) a network that is maintained across the inactive and the active states and (ii) a network that rearranges upon activation. Comparative analysis of GPCR crystal structures independently confirms the striking conservation of water-mediated interaction networks. These conserved water-mediated interactions near the G protein-coupling region, along with diverse water-mediated interactions with extracellular ligands, have direct implications for structure-based drug design and GPCR engineering.

## Significance

G protein-coupled receptors (GPCRs) represent both the largest class of drug targets and the largest family of human membrane proteins. Recent three-dimensional structures are revealing the presence of many water molecules buried inside GPCRs, but the functional role of these waters has been unclear. Using extensive atomic-level computer simulations, we find that although most of these waters are highly mobile, a few are stable. These stable water molecules mediate state-dependent and state-independent polar networks that are conserved across diverse GPCRs. These findings promise to help guide the rational design of GPCR-targeted drugs.

## Introduction

G protein-coupled receptors (GPCRs) are membrane proteins that act as a control panel for our cells: they recognize the presence of extracellular molecules such as hormones or neurotransmitters and stimulate intracellular signaling pathways in response. Humans have over 800 GPCRs, which share a common architecture comprising seven transmembrane (TM) helices (1).

GPCRs transmit signals across the cell membrane by transitioning from an inactive conformational state to an active conformational state. The active conformational state, which is favored by binding of certain ligands to the extracellular side of the GPCR, couples to and stimulates G proteins. Available structures indicate that diverse GPCRs undergo similar conformational changes upon activation (2). However, these GPCRs have evolved to recognize dramatically different ligands: human GPCRs recognize molecules ranging from amines, nucleosides, lipids, and steroids to entire proteins, in most cases with high specificity. This ability to stimulate cellular pathways in response to the presence of specific extracellular ligands makes GPCRs excellent drug targets, and in fact roughly one-third of all drugs act by binding to GPCRs (3). The process by which diverse ligands bind to GPCRs and favor activation is of great interest in molecular biology and drug discovery but remains incompletely understood.

Recent high-resolution crystal structures reveal dozens of water molecules buried in the transmembrane region of GPCRs (4–9). Water has been shown to play a critical role both in the binding of drugs to their targets (10) and in the function of many proteins (11, 12), raising the question of how the water molecules in GPCRs behave and what function they might serve. Indeed, this question was raised even before the high-resolution structures were available (13, 14).

Unfortunately, answering this question from crystal structures alone is difficult, because each crystal structure represents a static snapshot. In reality, water molecules may be highly mobile, and GPCRs themselves are constantly changing conformation (15). Moreover, most of the available GPCR crystal structures were solved at cryogenic temperatures at which water molecules are expected to be much less mobile than at body temperature. Even recovering average positions of water molecules from crystallographic data can be difficult when the corresponding electron density is not highly and unambiguously localized.

Here, we investigate the structural dynamics of water molecules in a diverse set of GPCRs using molecular dynamics simulations, which allow us to follow the trajectory of each individual water molecule at physiological temperature and in a non-crystallographic membrane environment. We compare water dynamics not only between different GPCRs but also between the active and inactive conformational states of the same GPCR. Our results show that the water molecules observed in GPCR crystal structures are not all equal: a few are stable in their crystallographically observed position, but most are highly mobile, spending only a small fraction of their time near that position.

Remarkably, stable water molecules near the G-protein-binding site form a network of polar interactions (hydrogen bonds) that rearranges upon receptor activation. This network is almost perfectly conserved across the GPCRs we examined, in sharp contrast to water-mediated interactions with ligands, which vary widely from one GPCR to another. This conserved water network appears to play a key role in the GPCR activation mechanism that was preserved as GPCRs evolved to recognize different ligands.

## Results

We focus on class A GPCRs, which represent both the largest class of GPCRs and the best characterized class structurally. We chose functionally distinct and evolutionarily distant ligand-activated class A GPCRs for which both inactive-state and active-state structures are available: β_2_ adrenergic receptor (β_2_AR (16, 17)), M2 muscarinic receptor (M2R (18, 19)), μ opioid receptor (MOR (20, 21)), and adenosine A_2A_ receptor (A_2A_R (22, 23)). We also included δ opioid receptor (DOR) as it has a particularly high-resolution inactive-state crystal structure (1.8 Å) (24). Collectively, these GPCRs bind to different types of ligands (amines, peptides, nucleosides) and different G proteins (Gs, Gi). All simulations included an explicitly represented lipid bilayer, water, salt ions, and the co-crystallized ligand (**Table S1**).

### Only a minority of crystallographic water molecules are stable

In simulations at physiological temperature, only a few of the waters resolved in high-resolution GPCR crystal structures remain near their crystallographically observed positions (**Fig. 1a** and **Fig. S1**). Our simulations are consistent with crystallographic data in that the same regions of the receptor are populated by waters in both cases (**Fig. S1**), but in simulation, most of the waters swim about, showing little preference for the specific positions occupied by waters in the crystal structures. A few crystallographic water positions, however, are consistently occupied by water molecules in simulation. We refer to these crystallographic water molecules as “stable,” even though an individual water molecule at such a position is generally replaced by a different water molecule at the same position at least every few hundred nanoseconds. In DOR simulations, for example, the water in the “kink” of transmembrane helix 6 (TM6) was highly stable with a water molecule observed within 1 Å of the crystallographic position for 99% of the time. On the other hand, the waters in the receptor core were highly mobile; for some of these, a water molecule was found within 1 Å of the crystallographic position less than 10% of the time.

**Fig. 1.**
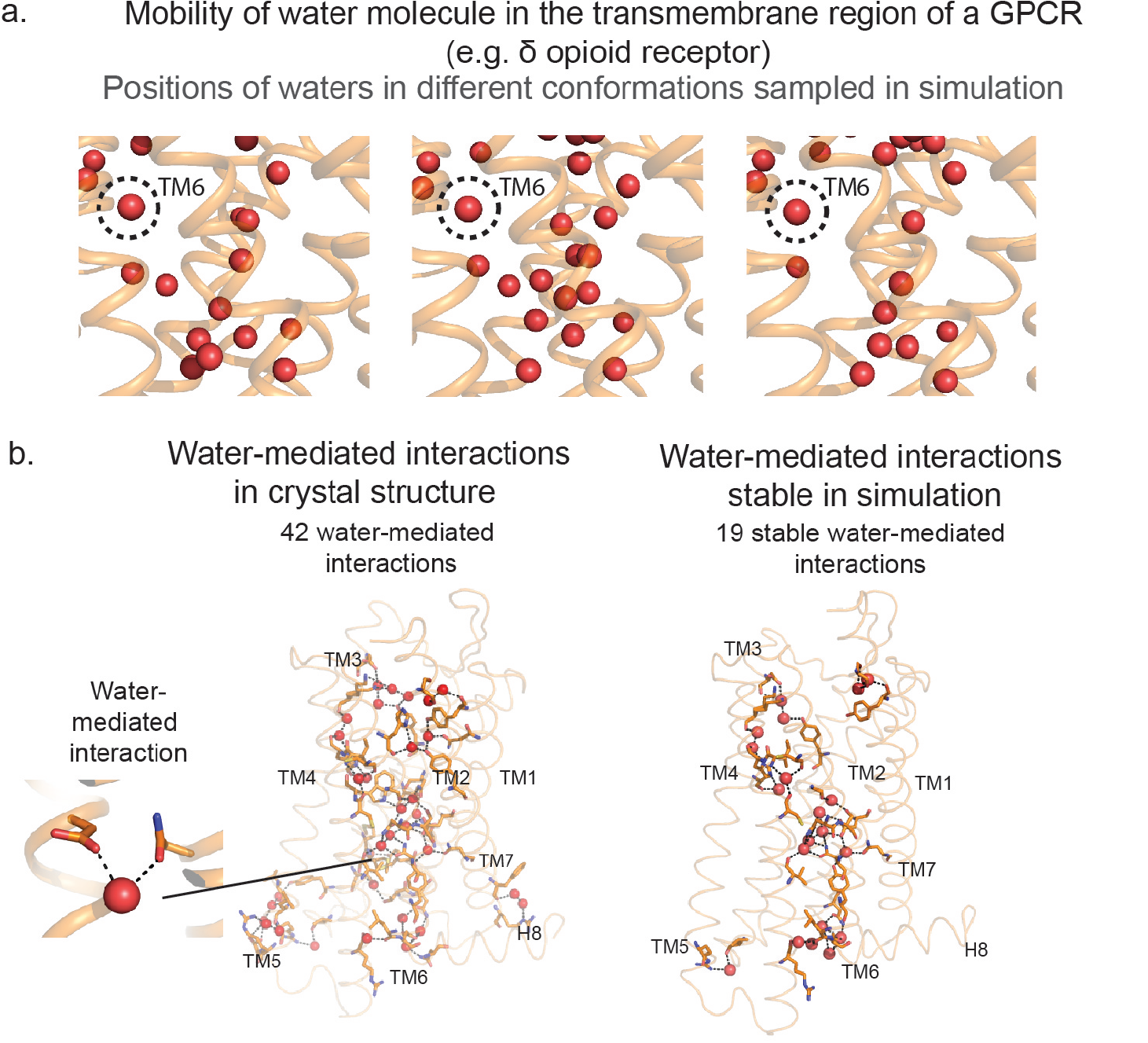
Stability of water molecules and water-mediated interactions in GPCRs. (a) Positions of water molecules (red spheres) in one region of a GPCR (the δ opioid receptor [DOR]) in three simulation snapshots. Receptor transmembrane helices are shown as orange ribbons. Most waters are mobile, but the water at the TM6 kink (circled) is not. (b) Left: water-mediated interactions present in the crystal structure of DOR. Right: water-mediated interactions formed over 60% of the time in simulation. Hydrogen bonds forming water-mediated interactions are shown as black dashed lines, and the residues involved in these interactions are shown with orange sticks.

To further characterize the behavior of water molecules in the structural framework of the receptor, we investigated the polar interactions (hydrogen bonds) in the receptor formed by the water molecules (**Fig. 1b**). We focused on water-mediated interactions in which a pair of receptor residues is connected by one or two water molecules in the network of hydrogen bonds. We estimated the stability of a given water-mediated interaction based on the frequency of the interaction’s presence in simulation (see **Methods**).

Only a fraction of the crystallographically observed water-mediated interactions are stable in simulation, because most of the water molecules are highly mobile, forming many transient, alternative polar interactions. For example, of the 42 water-mediated interactions in the DOR crystal structure, only 19 are formed in 60% or more of simulation frames (**Fig. 1b** and **Fig. S2**).

### Conserved networks of state-independent and state-dependent water-mediated interactions in GPCR activation

Might water molecules play a role in GPCR activation? To address this question, we first compared water-mediated interactions in simulations of inactive and active GPCR states. We also compared the water-mediated interactions across different GPCRs (β_2_AR, A_2A_R, MOR, and M2R). For the comparisons, we analyzed interactions between residues in structurally equivalent positions at different receptors. Structurally equivalent residues were identified using GPCRdb numbers (25), which are receptor-independent generic numbers for referencing structurally equivalent positions in GPCRs (see **Methods**).

We found that a surprisingly large set of stable water-mediated interactions (i.e., those formed over 60% of the time in simulation) were shared across *all* receptors examined in a given activation state (**Fig. 2**): six interactions mediated generally by four waters in the inactive state, and 12 interactions mediated generally by five waters in the active state.

**Fig. 2.**
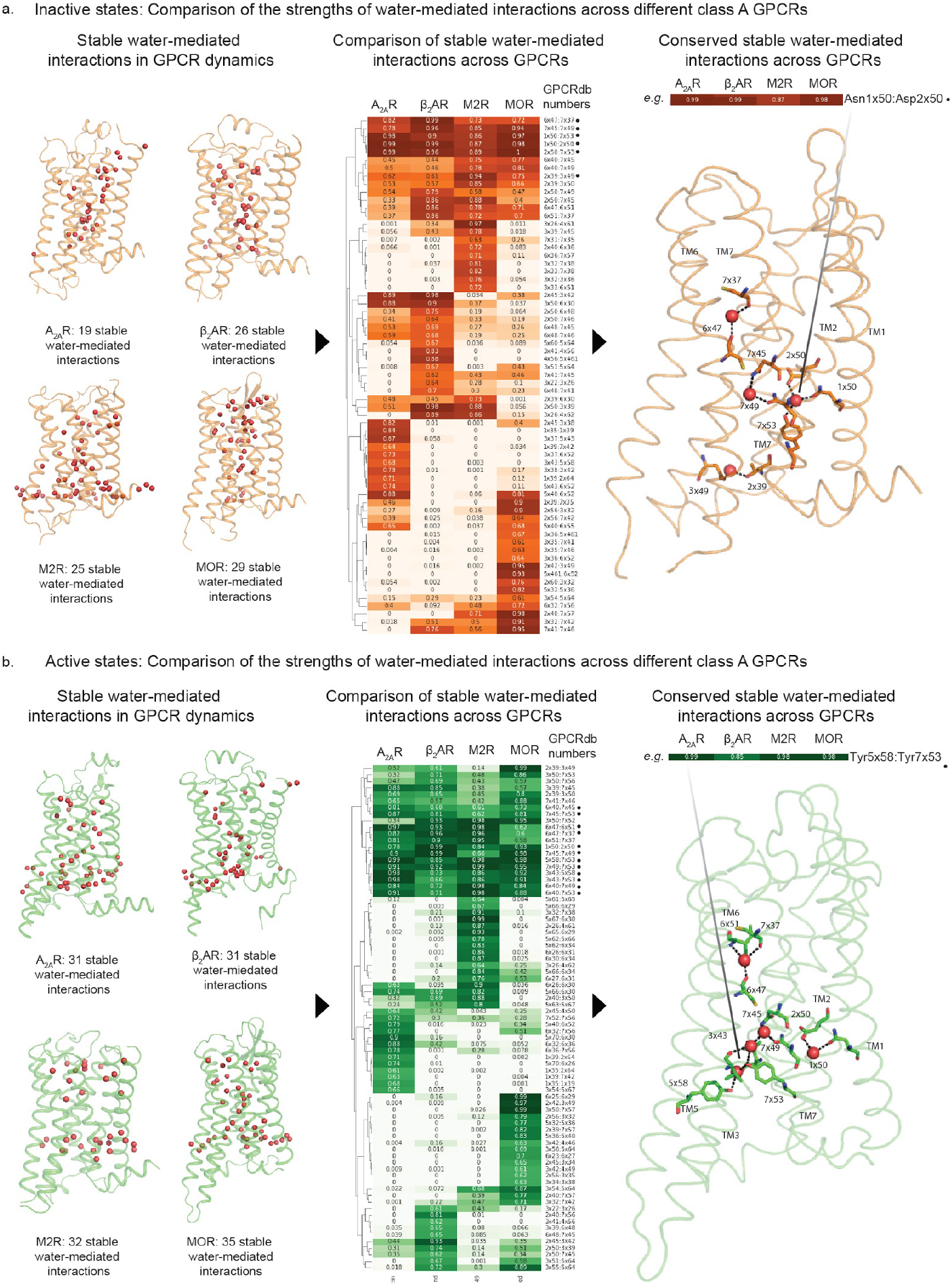
Conserved water-mediated interactions in inactive and active states of GPCRs. Comparison of the stability of structurally equivalent water-mediated interactions across different GPCRs in (a) inactive states and (b) active states. Four functionally distinct and evolutionarily distant class A GPCRs were compared: A_2A_R, β_2_AR, M2R, MOR The stability of the water-mediated interactions is estimated based on the frequency of formation in simulations. (a-b) Left: Frames from simulations of four functionally distinct GPCRs. Receptors are shown as ribbons and water molecules are shown as spheres. A schematic slider is shown to indicate that the each image represents a simulation frame. Middle: Heatmap comparing the stability of water-mediated interactions. Columns indicate receptors and rows indicate pairs of structurally equivalent positions, shown using GPCRdb numbers. The cells indicate the measure of stability of water-mediated interactions. Stability is shown as percentage values and using corresponding colors. Lighter shades indicate water-mediated interactions that have low stability and darker shades indicate water-mediated interactions that have high stability. Rows with dark cells in all columns indicate interactions that have highly stability in all the receptors being compared (marked with black dot). Right: Water-mediated interactions that are stable (frequency > 60%) across all four receptors mapped onto a GPCR structure. Hydrogen bonds forming water-mediated interactions are shown as black dashed lines, with the residues involved shown as sticks. Inactive state is shown in orange and active state is shown in green.

These conserved interactions represent 20−30% of the stable water-mediated interactions observed in each receptor individually. In contrast, a majority of the remaining stable interactions are found in just one of A_2A_R, β_2_AR, M2R and MOR. For example, of 32 stable water-mediated interactions in active-state M2R, 12 are shared across all four active-state receptors, and 14 of the remaining 20 are unique to M2R.

Among the conserved, stable, water-mediated interactions, three interactions are maintained in a state-independent manner, *i.e*. they are present in both the inactive states and the active states of A_2A_R, β_2_AR, M2R and MOR. While one of these interactions stabilizes a helical kink of TM6, the other two water molecules are proximal to the G protein-coupling region and mediate interactions between residues on TM1 and TM2 and between residues on TM7 (**Fig. 3a**).

**Fig. 3.**
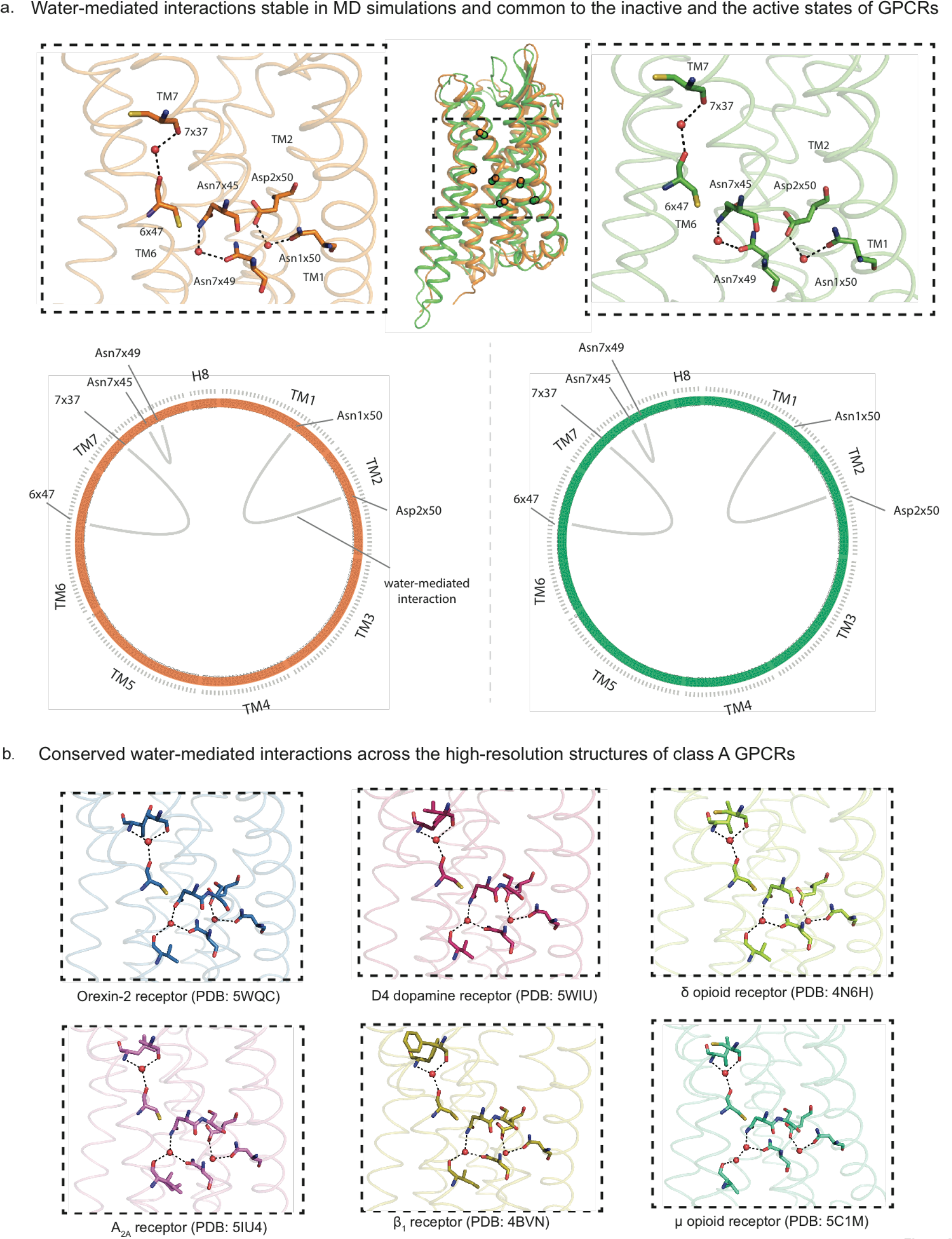
State-independent water-mediated interaction network: conserved and stable water-mediated interactions maintained across the inactive and the active states of diverse GPCRs. (a) Stable water-mediated interactions (frequency > 60%) present in both the inactive (orange) and the active (green) states of diverse GPCRs. Top: the conserved stable water-mediated interactions mapped onto a structure. Residues are shown as sticks, water molecules are shown as spheres, and hydrogen bonds forming water-mediated interactions are shown as black dashed lines. Bottom: The conserved stable water-mediated interactions are shown using ‘flareplots’. In flareplots, the amino acid residues in the transmembrane helices and helix 8 are shown as points on the circle. The water-mediated interactions between the residues are shown as chords connecting the points on the circle. (b) Water-mediated interactions common across the high-resolution structures (resolution of 2.1 Å or lower) of diverse GPCRs: inactive Orexin-2 (PDB ID: 5WQC), D4 dopamine (PDB ID: 5WIU), δ-opioid (PDB ID: 4N6H), A_2A_ adenosine (PDB ID: 5IU4) and β_1_ adrenergic (PDB ID: 4BVN) receptors, and active μ-opioid receptor (PDB ID: 5C1M). Residues are shown as sticks, water molecules are shown as spheres, and hydrogen bonds forming water-mediated interactions are shown as black dashed lines.

Two additional surprising observations link the conserved, stable, water-mediated interactions to the GPCR activation process. First, the remaining conserved, stable water-mediated interactions are in the lower half of the GPCR, near the G protein-binding interface, where the largest structural rearrangements occur upon activation. Second, although these interactions are conserved across diverse GPCRs in a given activation state, they differ dramatically between the active and inactive states. Every conserved stable water molecule near the G protein-binding interface changes interaction partners upon activation, and a number of water molecules form conserved, stable water-mediated interactions in the active state but not in the inactive state. Water-mediated interactions between TMs 1, 2, and 7 are stably formed in the inactive states (**Fig. 4a**). This network is disrupted upon activation and a new stable network of water-mediated interactions is formed between TM3, TM5, TM6, and TM7 (**Fig. 4a**). Tyr7×53 (GPCRdb number) of the NPxxY motif acts as a switch between the inactive and active water-mediated networks. In the inactive state, Tyr7×53 interacts with a water molecule at the TM1-TM2 interface, and upon activation Tyr7×53 switches conformation and interacts with water molecules at the interface of TMs 3, 5, and 6.

**Fig. 4.**
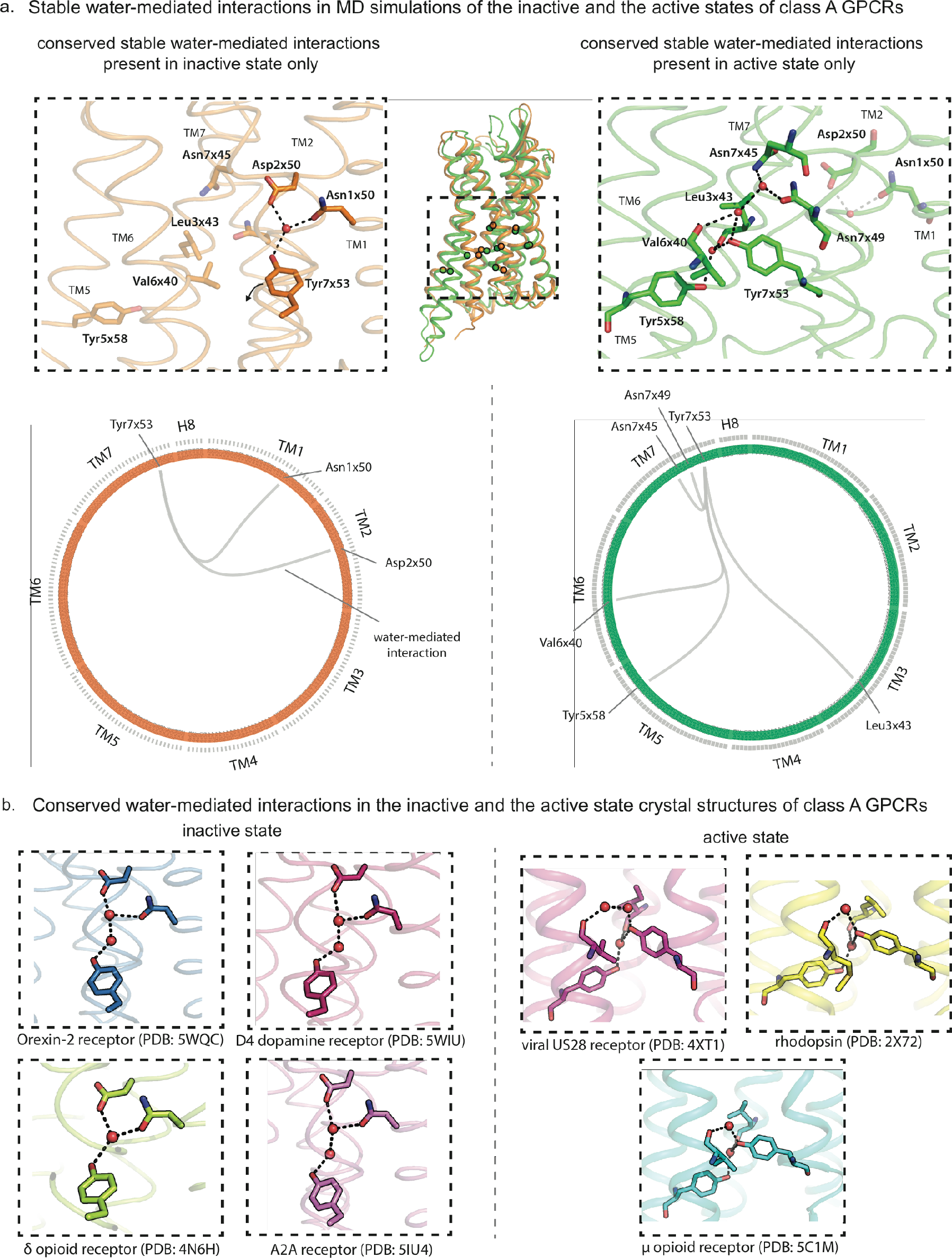
State-dependent water-mediated interaction network: conserved and stable water-mediated interactions maintained exclusively in the inactive states or the active states of diverse GPCRs. (a) Stable water-mediated interactions (frequency > 60%) present exclusively in all the inactive (orange) or all the active (green) states of diverse GPCRs. Top: the conserved stable water-mediated interactions of inactive state and active states mapped onto structure. Residues are shown as sticks, water molecules are shown as spheres, and hydrogen bonds forming water-mediated interactions are shown as black dashed lines. Bottom: The conserved stable water-mediated interactions are shown using ‘flareplots’. (b) Water-mediated interactions present exclusively in the crystal structures GPCRs in inactive state or active state. The high-resolution inactive state structures are of the following GPCRs: Orexin-2 (PDB ID: 5WQC), D4 dopamine (PDB ID: 5WIU), δ-opioid (PDB ID: 4N6H) and A_2A_ adenosine (PDB ID: 5IU4) receptors. The active state crystal structures of the following GPCRs are considered: rhodopsin (PDB ID: 2×72), viral chemokine receptor US28 (PDB ID: 4XT1) and μ-opioid receptor (PDB ID: 5C1M). Residues are shown as sticks, water molecules are shown as spheres, and hydrogen bonds forming water-mediated interactions are shown as black dashed lines.

All the side-chains involved in this rewiring of interactions with stable waters upon activation are highly conserved as polar residues across class A GPCRs (>89%) (**Table S2**). Taken together, our observations indicate that stable waters near the G protein-coupling interface form a conserved polar interaction network that rearranges upon receptor activation. The fact that this water-mediated activation switch was preserved as the GPCR family evolved suggests that it is functionally important.

### Comparative analysis of crystal structures confirms conservation of state-dependent and state-independent water-mediated interactions

We postulated that the conserved stable water-mediated interactions identified from the MD simulations would correspond to water-mediated interactions that are maintained across crystal structures of different receptors. In order to investigate this, we compared the water-mediated interactions from all available high-resolution crystal structures (see **Methods**): inactive Orexin-2, D4 dopamine, δ-opioid, A_2A_ adenosine and β_1_ adrenergic receptors, and active μ-opioid receptor (**Fig. 3b**). We compared the water-mediated interactions across structurally equivalent positions (see **Methods**). A network of seven water-mediated interactions involving three water molecules is maintained in all these crystal structures (**Fig. 3b**). Interestingly, this network of water-mediated interactions is highly similar to the network of conserved, state-independent, stable interactions observed in the MD simulations. All the three water-mediated interactions found to be conserved and stable across the simulations (**Fig. 3a**) are maintained consistently across the crystal structures (**Fig. 3b**). The remaining interactions that are consistently maintained in the crystal structures involve the same three waters as the conserved stable water-mediated interactions in the simulations.

Similarly, we compared the water-mediated interactions across high-resolution crystal structures of inactive states and crystal structures of active states. We found a network of two water-mediated interactions formed by Tyr7×53 to be maintained exclusively in the inactive state across Orexin-2, D4 dopamine, δ-opioid and adenosine A_2A_ receptors (**Fig. 4b**). The only exception where this network is absent is in the case of inactive β_1_ adrenergic receptor, and this is explained by the presence of a Tyr7×53Leu thermostabilizing mutation, which would prevent formation of any polar interactions. Interestingly, the conserved inactive state-dependent network identified from crystal structures is identical to the conserved, stable water-mediated interaction network found in the MD simulations of inactive states (**Fig. 4a,b**). Similarly, we also found a network of three water-mediated interactions formed by Tyr7×53 to be maintained exclusively in the crystal structures of active state rhodopsin, chemokine receptor US28, and μ-opioid receptor (**Fig. 4b**). All the three water-mediated interactions identified from the crystal structures of active states are present in the network of conserved, stable water-mediated interactions found in the MD simulations of the active states (**Fig. 4a,b**).

Taken together, the comparative analysis of crystal structures supports the presence of conserved state-dependent and state-independent water-mediated interactions in GPCRs.

### Stable water molecules in the ligand-binding pocket vary between different GPCRs

A number of the water molecules in high-resolution crystal structures of GPCRs form direct interactions with the ligand, and previous studies have pointed out the importance of water in determining ligand affinities and binding rates (26). We thus also examined the occupancy of waters in the binding pocket and the stability of water-mediated interactions between ligands and receptor residues in simulation.

Most waters in the binding pocket are swimming about. Certain waters and water-mediated interactions, however, are stable. For example, in simulations of inactive-state DOR, two waters in the binding pocket are highly stable near their crystallographic positions (**Fig. 5a**). These two stable waters mediate interactions between a hydroxyl group on the ligand and polar residues on TMs 5 and 6.

**Fig. 5.**
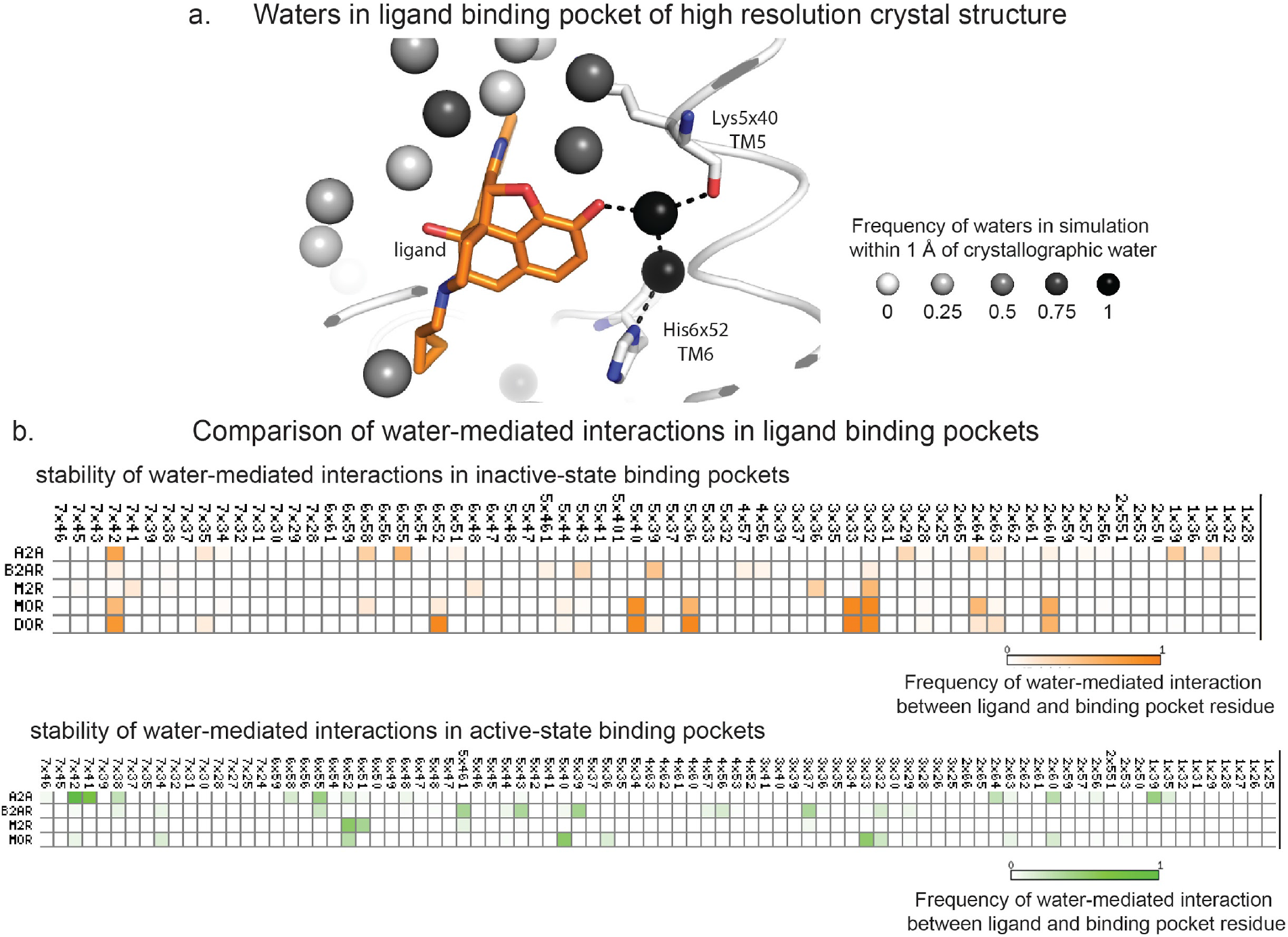
Diversity of stable water-mediated interactions in ligand-binding pockets of GPCRs. (a) Water molecules in the ligand-binding pocket of a high-resolution GPCR structure (DOR). The water occupancy in simulation of each crystallographic water position is shown on a white (0% occupancy) to black (100% occupancy) spectrum. Water occupancy in simulation is calculated as the fraction of simulation frames in which a water molecule is present within 1 Å of each crystallographic water positions. Two highly stable waters and the water-mediated interactions they form in the binding pocket are highlighted. The hydrogen bonds formed by these highly stable water molecules connecting the ligand and the binding pocket are shown as black dashed lines. (b) Heatmaps comparing the stability of water-mediated interactions in the ligand-binding pocket of five class A GPCRs: A_2A_R, β_2_AR, M2R, MOR, and DOR. In the heatmaps, the rows indicate receptors and the columns indicate structurally equivalent binding pocket positions. The frequency of water-mediated interactions between the ligand and the binding pocket residues in MD simulations is mapped to a spectrum of white to orange (for inactive state) and white to green (for active state). Lighter shades indicate water-mediated interactions that have low stability and darker shades indicate water-mediated interactions that have high stability.

No stable water-mediated interactions between the ligand and the receptor appear to be conserved across diverse GPCRs. Our simulations showed no such interactions conserved across all the receptors examined, and only very limited structurally equivalent interactions between subsets of these receptors (**Fig. 5b**). This variability in water-mediated interactions across ligand-binding pockets suggests that, unlike the G protein-coupling region, the binding pocket has been adapted to exploit different water-mediated interactions in different GPCRs as the GPCR family evolved.

## Discussion

The human genome includes hundreds of class A GPCRs, and these receptors have evolved to recognize extremely diverse extracellular ligands—ranging from nucleosides and amines to peptides and even entire proteins—often with very high specificity. On the other hand, all of these GPCRs couple to the same set of intracellular G proteins, which share very similar structures. One might thus imagine that GPCRs would have evolved a common activation mechanism. Indeed, previous studies have identified conserved networks of amino acid contacts and ionic interactions that appear to facilitate such a mechanism (2, 27).

Our results suggest that certain water molecules also play an important, conserved role in GPCR activation. In particular, we discovered a network of highly stable water molecules near the G protein-binding site that is conserved almost perfectly across diverse class A GPCRs and that rearranges in a conserved manner upon GPCR activation. Waters also play a critical role in mediating ligand binding in GPCRs, but the water-mediated interactions in the ligand-binding pocket vary widely among class A GPCRs, likely reflecting the fact that the binding pockets of these different GPCRs have evolved to recognize different ligands.

Our results complement those of previous experimental and computational studies. Recent high-resolution crystal structures have revealed the presence of dozens of waters within GPCRs. Analyses of earlier, lower-resolution crystal structures have suggested the presence of certain conserved structural water molecules (13, 14). Radiolytic footprinting has provided insights into tightly bound waters and dynamic waters in rhodopsin (28). Simulation-based studies have shown the presence of activation associated internal water pathways in GPCRs (28, 29) and indicated a role for waters in ligand binding (30–32).

A few caveats are in order. First, the force fields used for MD simulation are not perfect, although simulations have proven useful in elucidating water dynamics that are difficult to characterize experimentally (33, 34). Second, the strengths of hydrogen bonds in proteins can vary depending on factors such as the local structural environment (35), but our current analysis does not take this into account. Third, we lack a direct method to determine the functional importance of a water molecule in a particular position. We thus use stability and conservation across receptors as rough proxies for importance. Fourth, our analysis of stable water molecules and water-mediated interactions does not cover all the ways in which water may contribute to GPCR function; for example, displacement of certain water molecules from the binding pocket upon ligand binding is a determinant of ligand-binding affinity (36). Finally, our analysis has focused on class A GPCRs as they are the most extensively structurally characterized class of GPCRs. Future work will be necessary to understand the role of water molecules in the activation of other GPCR classes.

Our findings have several implications for GPCR drug discovery and GPCR engineering. First, understanding the behavior of water molecules in the binding pocket can aid structure-based drug discovery (26). Our results show that crystallographically resolved waters cannot all be treated as equal. One needs to characterize the dynamics of water molecules (*e.g*. through simulations) to identify the stable waters, which can then be exploited for drug design (37).

Second, the conserved stable waters and water-mediated interactions identified in this study can aid in structural modeling. Homology models of GPCRs are widely used for drug discovery, both when no crystal structure of the target is available and when the available structures are not in the desired activation state. The conserved water molecules identified in this study for active-state and inactive-state GPCRs can be incorporated into homology models and used to refine those models. Likewise, these water molecules can also be incorporated into low-resolution experimental GPCR structures, such as cryo-EM structures (38), in which few waters are resolved.

Third, mutating GPCR residues to stabilize either the active state or the inactive state has proven useful in GPCR crystallography as well as in functional investigations. The distinct water-mediated interaction networks associated with different activation states provide new avenues for biochemically perturbing the conformational landscape of GPCRs. In particular, residues interacting with the conserved, stable waters identified in this study are excellent candidate sites for introducing mutations to shift the conformational ensemble towards the active or inactive state.

More generally, our results reveal a “hidden” structural feature of GPCRs, a network of stable water-mediated interactions that is not evident from crystal structures alone. Our study also highlights the untapped potential of comparing structural dynamics across a drug target family to identify shared and distinct molecular features.

## Material and Methods

### MD simulation trajectories

The MD simulation trajectories of inactive β_2_AR, M2R and MOR and active β_2_AR were obtained from previously published studies (**Table S1**). New MD simulations were performed for inactive DOR and A_2A_R and active A_2A_R, M2R, and MOR (**Table S1**), as described below. The simulations were all performed using CHARMM force fields (**Table S1**). We verified that the simulations initiated from the inactive-state and active-state structures were maintained in their respective states.

### MD simulation system setup

The new simulations were initiated from the crystal structures from the Protein Data Bank (PDB) of inactive DOR (PDB ID: 4N6H), inactive A_2A_R (PDB ID: 5IU4), active A_2A_R (PDB ID: 5G53), active M2R (4MQS), and active MOR (5C1M). Prime (Schrödinger, Inc.) was used to model in missing side chains. Hydrogen atoms were added, and protein chain termini were capped with the neutral groups acetyl and methylamide. Residues Asp2×50 and Asp3×49 were protonated in the active-state simulations and deprotonated in the inactive-state simulations. All other titratable residues were left in their dominant protonation state at pH 7.0. The prepared protein structures were aligned on the transmembrane helices using the OPM database (39). For low-resolution structures, *i.e*. structures of active A_2A_R, and M2R, waters were added with Dowser(40), in addition to internal waters that were resolved in the crystal structure. Using Dabble (https://zenodo.org/badge/latestdoi/29268375), the structures were then inserted into a pre-equilibrated palmitoyl-oleoyl-phosphatidylcholine (POPC) bilayer, and solvated with 0.15 M NaCl in explicitly represented water.

### MD simulation force field parameters

We used the CHARMM36 parameter set for protein molecules, lipid molecules, and salt ions, and the CHARMM TIP3P model for water. Parameters for the co-crystallized ligands naltrindole (in inactive DOR), NECA (in active A_2A_R), BU72 (in active MOR), and iperoxo (in active M2R) were generated using the CHARMM General Force Field (CGenFF) (41) with the ParamChem server (paramchem.org), version 1.0.0. Parameters for the co-crystallized ligand ZM241385 (in inactive A_2A_R) were obtained from an earlier study (42).

### MD simulation protocol

Simulations were performed on GPUs using the CUDA version of PMEMD (Particle Mesh Ewald Molecular Dynamics) in Amber (43). Prepared systems were minimized, then equilibrated as follows: The system was heated using the Langevin thermostat from 0 to 100 K in the NVT ensemble over 12.5 ps with harmonic restraints of 10.0 kcal•mol^−1^•Å^−2^ on the non-hydrogen atoms of lipid, protein, and ligand, and initial velocities sampled from the Boltzman distribution. The system was then heated to 310 K over 125 ps in the NPT ensemble with semi-isotropic pressure coupling and a pressure of one bar. For all the simulations except active A_2A_R, further equilibration was performed at 310 K with harmonic restraints on the protein and ligand starting at 5.0 kcal•mol^−1^•Å^−2^ and reduced by 1.0 kcal•mol^−1^•Å^−2^ in a stepwise fashion every 2 ns, for a total of 10 ns of additional restrained equilibration. For active A_2A_R, further equilibration was performed at 310 K with harmonic restraints on the protein and ligand starting at 5.0 kcal•mol^−1^•Å^−2^ and reduced by 1.0 kcal•mol^−1^•Å^−2^ in a stepwise fashion every 2 ns to 1.0 kcal•mol^−1^•Å^−2^ and then by 0.1 kcal•mol^−1^•Å^−2^ in a stepwise fashion every 2 ns, for a total of 28 ns of additional restrained equilibration.

Multiple independent simulations were initialized from the final snapshot of the restrained equilibration. These simulations were conducted in the NPT ensemble at 310 K and 1 bar, using a Langevin thermostat and Monte Carlo barostat. Simulations used periodic boundary conditions. A time step of 4.0 fs with hydrogen mass repartitioning (44) was used for inactive DOR, inactive A_2A_R, active M2R, and active A_2A_R, and a time step of 2.5 fs was used for active MOR. The simulation frames of inactive DOR, inactive A_2A_R, active M2R, and active A_2A_R were written every 200 ps and those of active MOR were written every 100 ps.

The active-state simulations were maintained in the active state using harmonic restraints applied to receptor residues within 5 Å of the co-crystallized nanobody for active M2 receptor and active MOR and within 5 Å of the mini G protein for active A_2A_R receptor. For active M2 receptor and active A_2A_R receptor, the harmonic restraints were 5.0 kcal•mol^−1^•Å^−2^. For active MOR, the harmonic restraints were 1.0 kcal•mol^−1^•Å^−2^.

Bond lengths to hydrogen atoms were constrained using SHAKE. Non-bonded interactions were cut off at 9.0 Å, and long-range electrostatic interactions were computed using the particle mesh Ewald (PME) method with an Ewald coefficient β of approximately 0.31 Å and B-spline interpolation of order 4. The FFT grid size was chosen such that the width of a grid cell was approximately 1 Å.

### Computation of water-mediated interactions

Hydrogen bonds in the MD simulations were computed using the HBonds plugin in VMD (45) with the following geometric criteria: donor to acceptor distance less than 3.5 Å and donor-hydrogen-acceptor angle greater than 110°. Hydrogen bonds in the high-resolution crystal structures were computed based on the donor to acceptor distance of 3.2 Å only, as crystal structures generally lack hydrogens. For the crystal structure-based analysis, we considered high-resolution structures (of resolution 2.1 Å or lower) of diverse GPCRs in inactive state: Orexin-2 (PDB ID: 5WQC), D4 dopamine (PDB ID: 5WIU), δ-opioid (PDB ID: 4N6H), A_2A_ adenosine (PDB ID: 5IU4) and β_1_ adrenergic (PDB ID: 4BVN) receptors. For the active state, crystal structures of the following GPCRs were considered: rhodopsin (PDB ID: 2×72), viral chemokine receptor US28 (PDB ID: 4XT1) and μ-opioid receptor (PDB ID: 5C1M). A water-mediated interaction between a pair of residues is defined when both residues form hydrogen bonds with the same water molecule or a pair of water molecules that in turn are linked by a hydrogen bond. In the ligand-associated analyses, water-mediated interactions are defined to occur when a water molecule or pair of water molecules connect a ligand atom and a residue through hydrogen bonds. We only considered interactions mediated by residues that were at least 4 positions apart in TM1-7 and helix 8 (H8) in order to avoid local short range interactions. We evaluated the presence of water-mediated interactions between structurally equivalent residues across different GPCR structures. Structural equivalence was assigned using the GPCRdb numbering scheme (25), which is developed based on the Ballesteros Weinstein numbering scheme (46) and corrects for helical bulges and constrictions. In the GPCRdb numbering scheme, every residue is addressed using two numbers separated by an ‘x’. The first number denotes the helix (1–8) and the second number denotes the residue position relative to the most conserved position on that helix, which is assigned the number 50. For example, 3×51 denotes a residue in transmembrane helix 6, one position after the most conserved residue (3×50). A bulge residue is assigned the same number as the preceding residue followed by a 1, *e.g*. 551 for a bulge following position 55.

### Computing frequencies of water-mediated interactions

Water-mediated interactions were computed for every frame of the MD simulations. The stability of a water-mediated interaction between a pair of residues was defined as the fraction of frames in which either a direct or extended water-mediated interaction is formed. The stability of water-mediated interactions was computed for various receptors in both the inactive and active states (**Table S1**).

### Water occupancy and water density analysis

To compute the occupancy of water molecules proximal to each crystallographic water position, we identified amino acid residues within 5 Å of the crystallographic water’s position and aligned all simulation frames to these residues. We then calculated the fraction of simulation frames in which a water molecule is within 1 Å of the crystallographic water position.

To computing water density, a grid with 1 Å^3^ voxels was superimposed upon a GPCR structure. We then computed the fraction of simulation frames in which an oxygen atom of a water molecule is contained within each voxel. The water density map was output as a ‘.dx’ file and loaded into VMD with an ‘isosurface’ representation to render a water density map.

### Sequence analysis

We obtained the human class A non-olfactory GPCR sequences from GPCRdb (25) and computed the percentage conservation of each amino acid at every position in TMs 1-7 and helix 8. The percentage of polar amino acids at each position was computed by adding the percentages of the following amino acids: Asn, Asp, Glu, Gln, His, Ser, Thr, and Tyr.

### Visualisation

Water-mediated interactions between residues were visualized using flareplots (https://gpcrviz.github.io/flareplot/). Structural visualization of protein structure and water-mediated interaction networks was performed using VMD (45) and PyMOL.

---

**Fig. S1.**
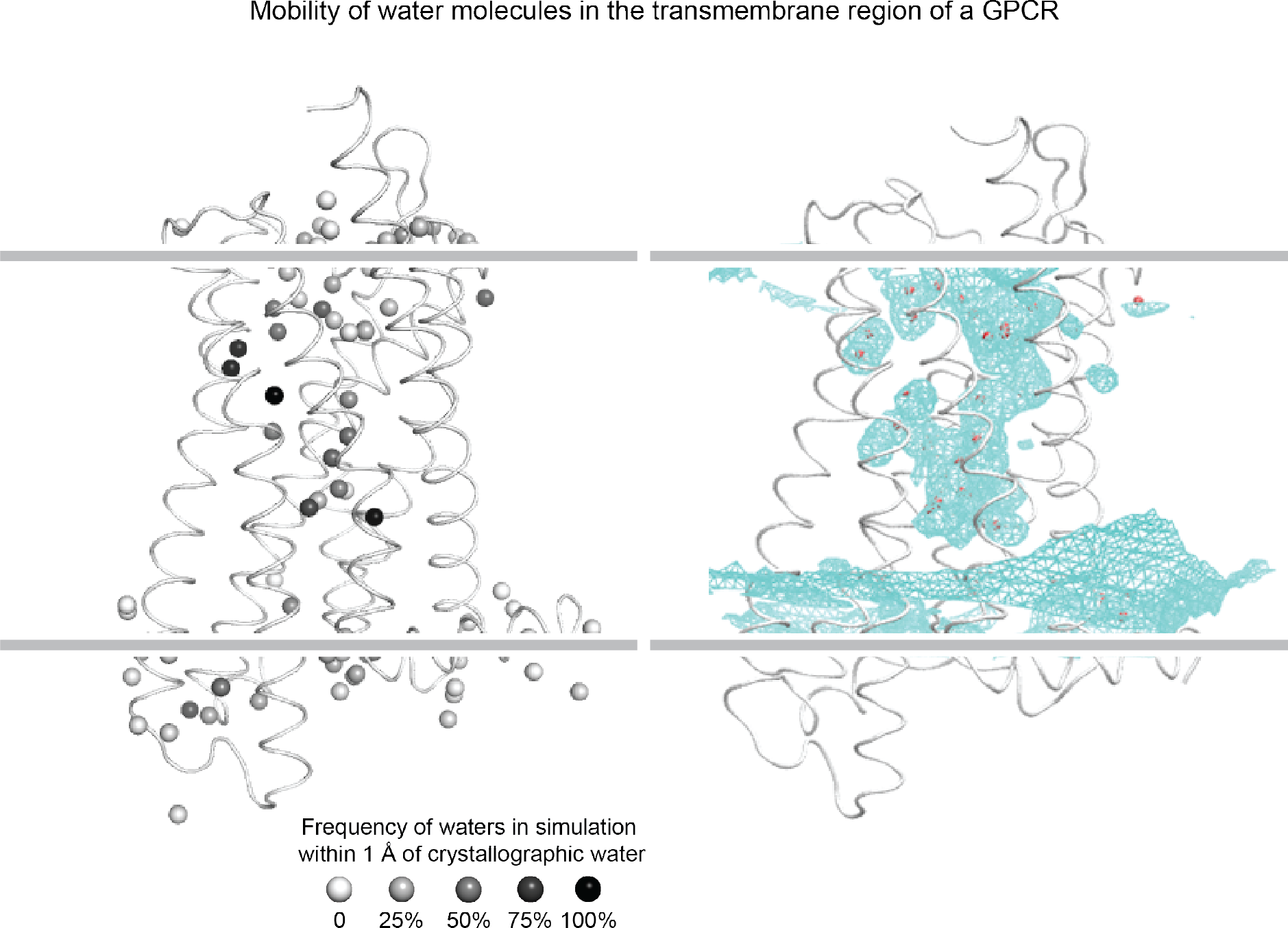
Occupancy of water molecules in GPCR simulation. (a) The occupancy of water molecules in simulations of DOR at crystallographic water positions. The occupancy of each position is shown on a white (0% occupancy) to black (100% occupancy) spectrum. Occupancy of waters in simulation is calculated as the fraction of simulation time where waters are present within 1 Å of each crystallographic water position. (b) The density of water molecules in the transmembrane region. The mesh encloses regions where waters in simulation are present at least 1% of simulation frames in each grid cell of dimension 1 Å × 1 Å × 1 Å. Crystallographic water positions are shown by red spheres. These almost always fall within the mesh-enclosed regions, indicating that despite the low occupancy of waters in simulation at most crystallographic water positions, waters tend to occupy the same regions of the receptor in crystal structures and simulations.

**Fig. S2.**
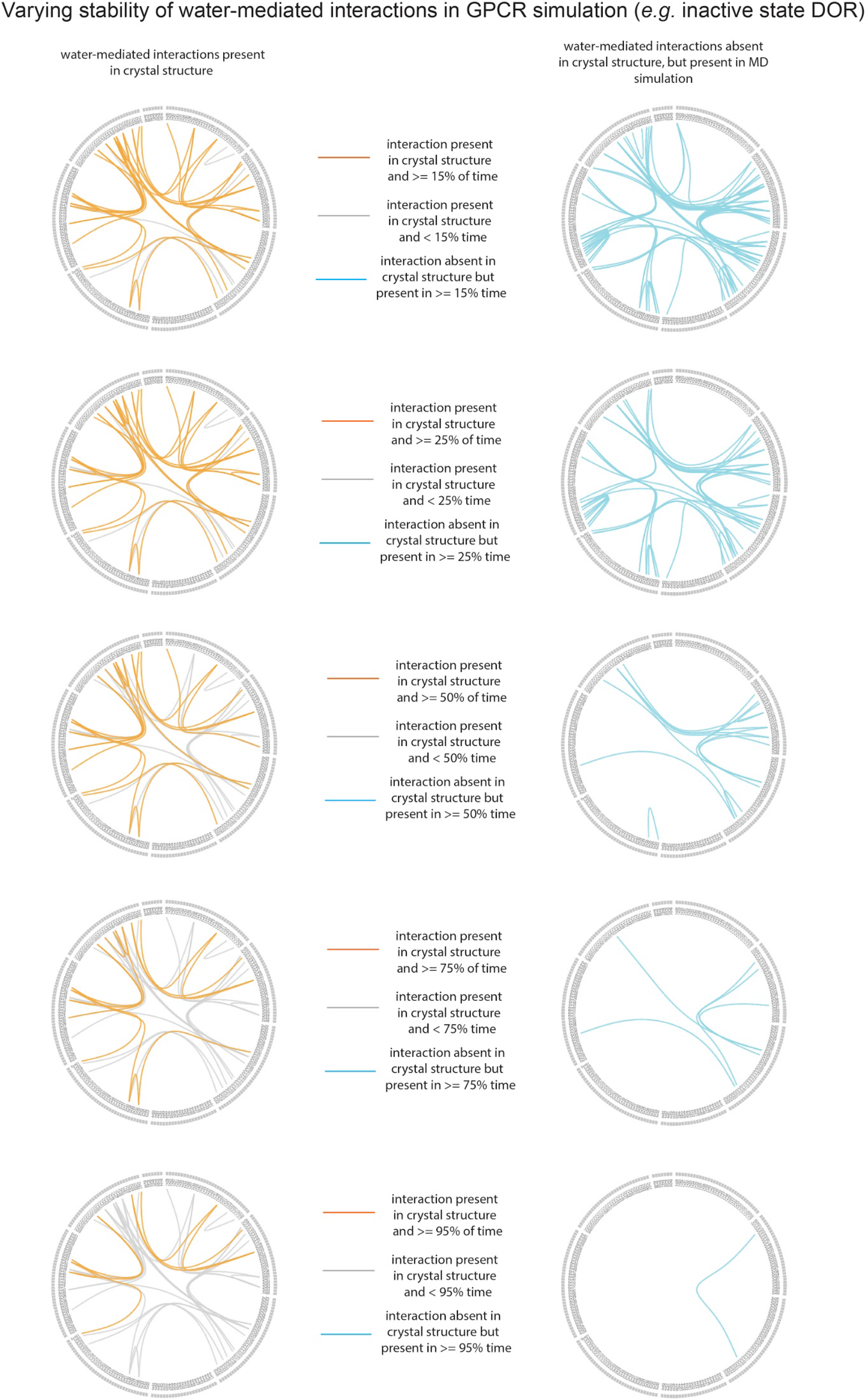
Stability of water-mediated interactions present in crystal structures and simulation. Water-mediated interactions present in MD simulation of a GPCR were stratified into three sets: interactions that are present in crystal structure and in simulations above a frequency threshold (orange), interactions that are present in crystal structure and in simulations below a frequency threshold (gray), interactions that are absent in crystal structure but are observed in simulation above a frequency threshold (gray). The water-mediated interactions were stratified at different frequency thresholds: 15%, 25%, 50%, 75%, and 95%.

**Supplementary Table S1.**
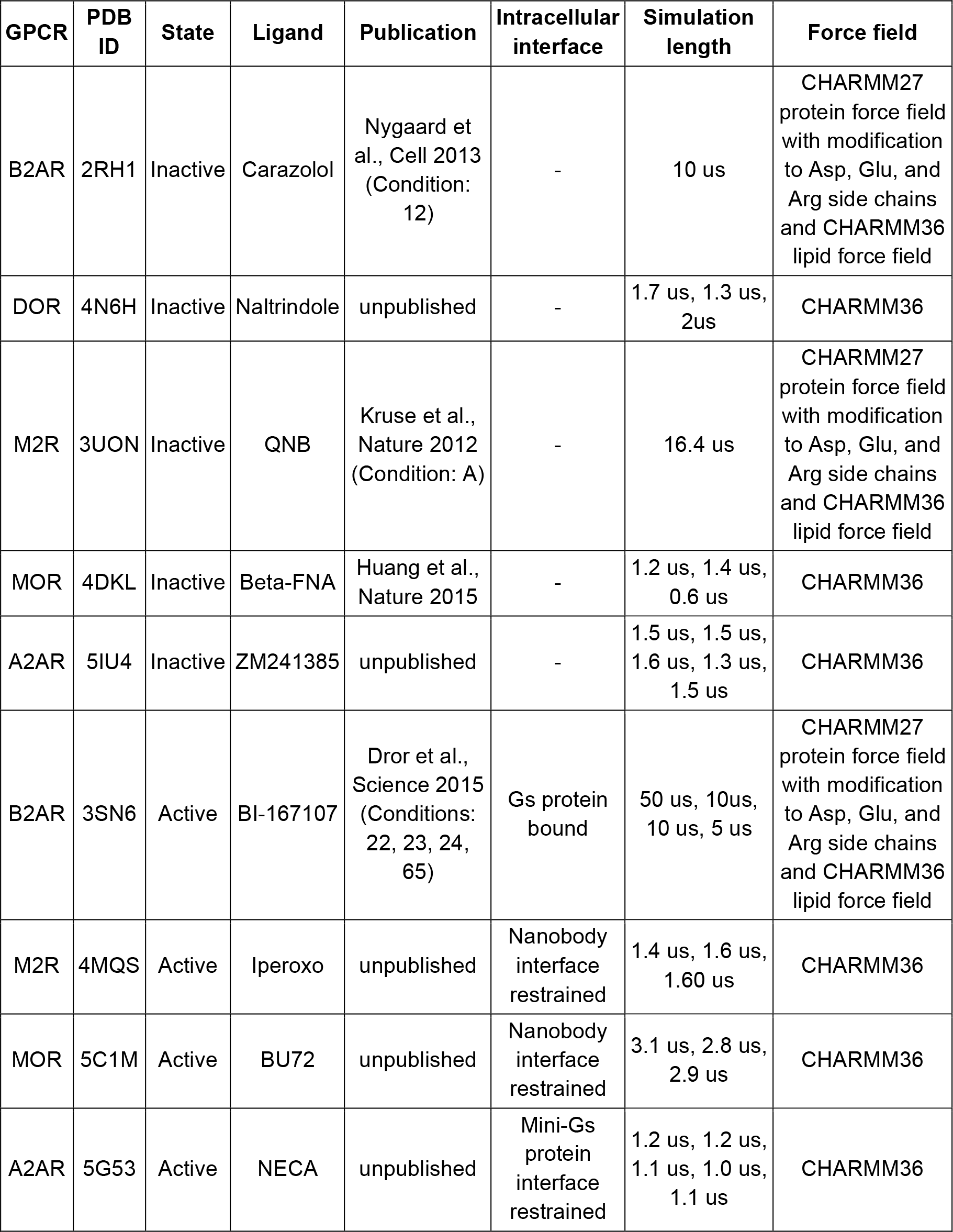
List of GPCR simulations analyzed in this study

**Supplementary Table S2.** Sequence conservation across human class A GPCRs (nonolfactory) of positions forming conserved stable water-mediated networks exclusively in inactive state and active state of GPCRs

| **GPCRDB**<br>**number** | **Polar**<br>**amino acids** | **Asn** | **Asp** | **Glu** | **Gln** | **His** | **Ser** | **Thr** | **Tyr** |
| --- | --- | --- | --- | --- | --- | --- | --- | --- | --- |
| 1x50 | 100% | 100% | 0% | 0% | 0% | 0% | 0% | 0% | 0% |
| 2x50 | 98% | 1% | 96% | 1% | 0% | 0% | 0% | 0% | 0% |
| 5x58 | 89% | 7% | 0% | 0% | 1% | 1% | 1% | 2% | 77% |
| 7x45 | 90% | 69% | 0% | 0% | 1% | 8% | 12% | 0% | 0% |
| 7x49 | 98% | 78% | 17% | 0% | 0% | 1% | 1% | 1% | 0% |
| 7x53 | 92% | 0% | 1% | 0% | 0% | 0% | 0% | 0% | 91% |

